# Identification of DOT1L Inhibitor in a Screen for Factors that Promote Dopaminergic Neuron Survival

**DOI:** 10.1101/2022.08.23.505021

**Authors:** Jun Cui, Joseph Carey, Renee A. Reijo Pera

## Abstract

Parkinson’s disease (PD) is a common neurodegenerative disorder characterized by the progressive loss of dopaminergic (DA) neurons in the substantia nigra region of the midbrain. Diagnostic criteria for PD require that at least two of three motor signs are observed: tremor, rigidity, and/or bradykinesia. The most common and effective treatment for PD is Levodopa (L-DOPA) which is readily converted to DA and has been the primary treatment since the 1960’s. Dopamine agonists have also been developed but are less effective than L-DOPA. Although the lack of a model system to study PD has hampered efforts to identify treatments, diverse screening strategies have been proposed for identification of new pharmaceutical candidates. Here, we describe a pilot screen to identify candidate molecules from a bioactive compound library, that might increase formation, maintenance and/or survival of DA neurons *in vitro.* The screen used a previously characterized reporter construct consisting of the luciferase gene inserted downstream of the endogenous tyrosine hydroxylase (TH) gene and neurons differentiated from human pluripotent stem cells for 18 days. The reporter mimics expression of TH and includes a secreted luciferase whose activity can be measured non-invasively over multiple timepoints. Screening of the bioactive compound library resulted in the identification of a single molecule, SGC0946, that is an inhibitor of DOT1L (Disruptor Of Telomeric silencing 1-Like) which encodes a widely-conserved histone H3K79 methyltransferase that is able to both activate and repress gene transcription. Our results indicate that SGC0946 increased reporter luciferase activity with a single treatment at 8-hours post-plating being equivalent to continuous treatment. Moreover, data suggested that the total number of neurons differentiated in the assays was comparable from experiment to experiment under different SGC0946 treatments over time. In contrast, data suggested that the survival and/or maintenance of DA neurons might be specifically enhanced by SGC0946 treatment. These results confirm other reports that indicate inhibition of DOT1L may play an important role in maintenance and survival of neural progenitor cells (NPCs) and their lineage-specific differentiation.

## Introduction

Parkinson’s disease (PD) is a common neurodegenerative disorder that affects almost one million people in the United States and more than 6 million internationally (Collaborators, 2018). Although the symptoms of PD vary among patients, a central hallmark is the progressive loss of movement control associated with the progressive loss of dopaminergic (DA) neurons of the substantia nigra of the midbrain (Albin et al., 1995). Intensive study over the years has identified numerous environmental and genetic risk factors that lead to PD in a subset of cases; however, the cause of the majority of PD remains poorly understood (Bloem et al., 2021).

One of the major limitations that hinders research in PD is the lack of access to affected human DA neurons. Thus, the development of human pluripotent cells including both human embryonic stem cells (hESCs) and induced pluripotent stem cells (iPSCs) has demonstrated the potential to provide a reliable source to obtain human DA neurons and study the disease. hESCs are pluripotent cells that have been derived from the inner cell mass of an early-stage embryo (Thomson et al., 1998) while iPSCs are generated from adult somatic cells, usually skin fibroblast cells, that are induced to pluripotency via a combination of four transcription factors (Takahashi and Yamanaka, 2006, Takahashi et al., 2007). Advances over the last decade or so, in defining differentiation conditions have made it possible to directly drive differentiation of human pluripotent cells to DA neurons and use these derived neurons to model PD in a dish (Yang et al., 2008, Swistowski et al., 2010, Gonzalez et al., 2013, Chambers et al., 2009, Zhang et al., 2014, Kriks et al., 2011, Kedariti et al., 2022, Stern et al., 2022, Singh et al., 2017, Barbuti et al., 2020). Multiple studies from different research laboratories have also shown that these derived neurons not only have cellular and biochemical characteristics very similar to the neurons developed in the human brain but also are functional when transplanted into animal models (Byers et al., 2015, Doi et al., 2014, Grealish et al., 2014, Kriks et al., 2011, Byers et al., 2011, Hoban et al., 2020) Thus, studies over the last decade plus have demonstrated that human pluripotent stem cells can provide a reliable source for generating DA neurons for *in vitro* disease modeling and have the potential to contribute to regenerative treatment options for the disease (Behl et al., 2022).

Developing treatments for PD traditionally has focused predominantly on relief of symptoms, as there is currently no effective mean(s) to restore DA neurons that are lost in PD patients. One potentially feasible option is to restore these neurons in patients via transplantation of *in vitro* derived DA neurons. Another approach is to screen for chemicals that have a protective effect directly on DA neurons. Pharmaceutical chemicals that may improve survival of DA neurons have multiple uses in PD related research and future applications: The immediate application of such chemicals is to enhance the production of a larger number of DA neurons more efficiently in *in vitro* derivation of DA neurons. Further, pharmaceutical chemicals would also provide a window into PD to enable mapping of biological function(s) of pathways that are compromised in PD. Finally, such chemicals might also slow or halt loss of DA neurons *in situ* during progression of PD.

As indicated above, DA neurons differentiated *in vitro* have the potential to provide high quality human midbrain DA neurons that are fully functional both in *in vitro* and *in vivo* assays and applications. Yet, it is clear that current differentiation methods are still time consuming and the number of derived neurons is often limited greatly by variation in the differentiation efficiencies from one experimental replicate to another. Applications that use derived DA neurons either for *in vitro* cellular modeling or for future cell replacement therapies would benefit greatly from the improvement of the differentiation methods and/or development of methods that distinguish DA neuron survival in a complex cell mixture. Here, we review the characterization of an hESC reporter line that has a secreted luciferase reporter gene inserted into the endogenous tyrosine hydroxylase (TH) gene in order to provide a rapid and efficient method to measure differentiation and survival of DA neurons in complex differentiation cultures (Bademci et al., 2012) Then, we present results of a pilot screening study, based on a well-characterized, commercially-available Tocris chemical library, that was aimed at identifying compounds that may improve the survival of DA neurons during direct differentiation *in vitro*, in adherence to culture plates and/or in post-differentiation survival.

## Materials and Methods

### The Tocris library

We chose to screen the Tocriscreen 2.0 library for potential factors that would act to promote differentiation, adherence and/or survival of DA neurons. This library was obtained from Bio-Techne Corporation (Minneapolis, MN) in a 96-well format with each chemical dissolved in 10mM DMSO with an indicated shelf-life of at least 6 months. The 1280 compounds within the Tocriscreen 2.0 library cover such research areas as cancer, immunology, neuroscience and stem cells. Pathways include the G-protein coupled receptors (GPCRs), kinases, enzymes and enzyme-linked receptors, chemicals that target pathways of cell biology and ion channels. For screening, day 18 DA neuron progenitors were plated and cultured for 10 days with DMSO only (control) or treated with chemicals during the first 48 hr (single treatment) or 8 days (continuous treatment). TH expression was detected using the luciferase assay and normalized to the treatment with DMSO only. Results were graphed with X axis representing the reading of single chemical treatment and Y axis representing continuous treatment for 8 days. Luciferase results were confirmed by Q-PCR of the luciferase and TH RNA.

### Directed dopaminergic differentiation and quantification of TH using luciferase assay

Pluripotent human stem cells (hESCs) were differentiated towards a dopamingeric neuron cell fate following the floor-plate induction method(Zhang et al., 2014). Cell supernatants were collected every day for the first 20 days of neuronal differentiation and every other day afterwards to ensure that the protocol produced DA neurons. Luminescent reactions were performed by mixing the supernatants with 10× reaction buffer consist of 300 mM Tris-HCl pH 8.0, 40 mM NaCl, 1% Triton X-100 and 200 μM substrate Coelenterazine (NanoLight; Flagstaff, AZ). Luminescent signal was measured using a Synergy H1 Plate Reader (BioTek; Santa Clara, CA). Background signal level was measured using blank medium with reaction buffer and subtracted from experimental data.

### Drug screening assays

hESCs were differentiated towards dopamingeric neurons for 18 days as described above. One parameter that directly affected the yield of DA neurons in the final culture in our previous study was cell density (Cui et al., 2016). To identify the optimal cell density for DA neuron differentiation and test performance of the TH luciferase reporter assay, we replated differentiated neural cells at different densities and followed the DA neuron growth for two weeks. When cells were plated at densities higher than 400 K cells per well, restraint by the surface growth area greatly limited neuron attachment and growth resulting in massive cell detachment within several days. In contrast, fewer than 50 K cells per well led to very poor cell attachment and a low total number of DA neurons. Within the optimal range, each well produced 10^3^ to 10^4^ units of luminescence signal when the culture matured and signals showed a linear correlation with cell numbers on Day 40, suggesting the cell growth was not restricted during the culturing period. The growth test was also performed in the 384-well format and exhibited a similar result with optimal cell densities of 20 K to 100 K cells per well. These results indicated that the reporter is suitable for used as luciferase based quantitative assays within the optimal range of cells per well. Given these results, on Day 18, neurons were dissociated by incubating with Accutase (Innovative Cell Technologies; San Diego, CA) for 5 minutes. Dissociated cells were counted and replated to 96-well plates coated with poly-L-ornithine/laminin/fibronectin at a density of 3 × 10^5^ cells/cm^2^ and allowed to recover for one week in neurobasal medium containing B-27 supplement, BDNF, GDNF, cAMP, TGFb, and ascorbic acid. Drug treatment was performed as previously described for neurotoxins (Grealish et al., 2014). Fleshly prepared Tocris solutions were diluted with the basal medium and added to each well to reach the desired final concentration. Assays were done with multiple wells of 4 replicates for each condition. Whole supernatants of the same well were collected before drug treatment and 48 hours post treatment for quantification using luciferase assay as described above. Viability of TH+ cells was calculated as the percentage of luminescent signal intensity post treatment compared to that before treatment of the same well.

### RNA extraction and quantitative PCR analysis

Total RNA was extracted using RNeasy mini kit (Qiagen, Hilden, Germany). For first-strand cDNA synthesis, 2 μg of total RNA was reverse transcribed with oligo-dT primers and SuperScript III reverse transcriptase (Life Technologies, Carlsbad, CA). Quantitative PCR reactions were run using Power SYBR PCR mix (Life Technologies, Carlsbad, CA) and detected by ABI 7300 Real-Time PCR System. Primers used for PCR are listed in **Table 1**. Relative quantity for each sample and gene was normalized and calculated based on the ΔCt values using GAPDH as the control.

**Table 1.**
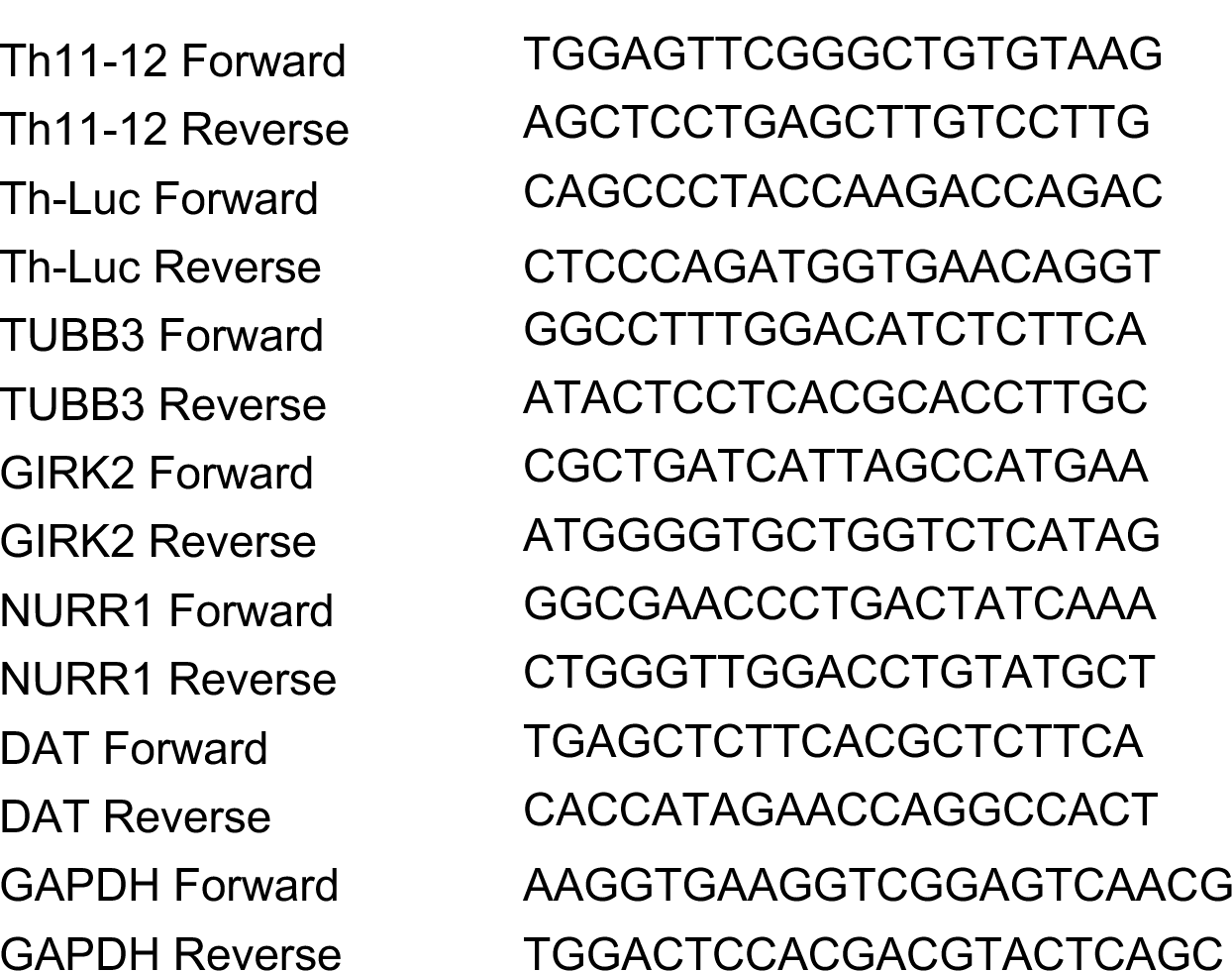
Primers Used in This Study.

## Results

In order to screen for chemicals that might enhance DA differentiation and/or survival, we used a genetic reporter system that we previously constructed and demonstrated was able to monitor differentiation efficiency and compare DA survival under different conditions (Cui et al., 2016). We note here that the reporter contains an insertion of a luciferase reporter gene into the endogenous TH locus, the gene that encodes the enzyme governing the rate-limiting step in dopamine production (Cui et al., 2016). The luciferase reporter was shown to enable rapid non-invasive quantification of dopaminergic neurons in cell culture throughout the entire differentiation process (Cui et al., 2016). Moreover, luciferase was shown to be under the same endogenous regulation of the TH gene demonstrating that the cellular assay is effective in assessing neuron response to different cytotoxic chemicals and able to be scaled for high throughput applications suggesting feasibility of use as a quantitative cellular model for toxin evaluation and drug discovery.

We differentiated the reporter-containing pluripotent hESCs, previously described (Cui et al., 2016) to midbrain DA neurons using an established floor-plate induction protocol with defined combinations of growth factors (Kriks et al., 2011, Zhang et al., 2014). The engineered cells had previously been shown to differentiate to TH+ DA neurons with similar robustness as the parental cell line in both a 96-well and 384-well format, with stable expression of TH observed starting from Day 11 of induction and continuing to increase throughout the following culturing, which correlated with the timing of midbrain DA neuron growth and maturation (Cui et al., 2016).

To screen the Tocriscreen 2.0 library, day 18 DA neuron progenitors were plated and cultured for 10 days with DMSO only (control) or treated with chemicals during the first 48 hr (single treatment) or 8 days (continuous treatment) as diagrammed **(Figure 1A)**. TH expression was detected using the luciferase assay and normalized to the treatment with DMSO only. Results were graphed with X axis representing the reading of single chemical treatment and Y axis representing continuous treatment for 8 days. We identified a single chemical component of the Tocriscreen 2.0 library, SGC0946, that resulted in a significant increase (>175% increase with either single or continual treatment relative to the DMSO Only control (**Figure 1B)**. We noted that multiple other chemical components of the Tocris library demonstrated smaller effects while the majority of chemicals demonstrated a reduction in luciferase activity relative to the DMSO Only control.

**Figure 1.**
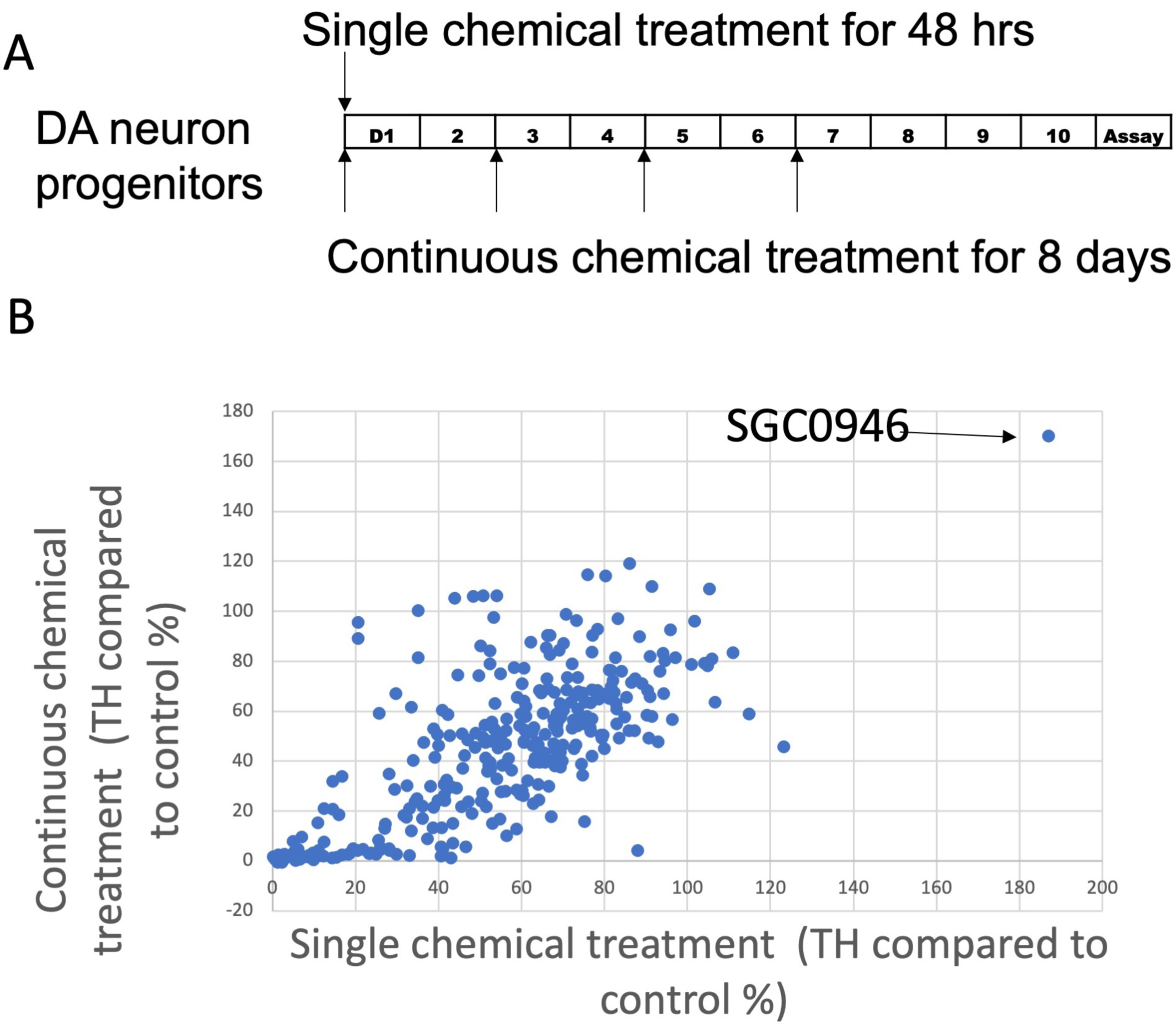
Screen of Tocriscreen 2 chemical library. **(A)** Experimental design. Day 18 DA neuron progenitors were plated and cultured for 10 days with DMSO only (Control) or treated with chemicals during the first 48 hr (single treatment) or 8 days (continuous treatment). Luciferase readings were done at the end of the 10-day differentiation. **(B)** Screen results. TH expression was detected using the luciferase assay and normalized to the treatment with DMSO only. X axis represents the reading of single chemical treatment and Y axis represents continuous treatment for 8 days.

We then compared TH expression as assessed by luciferase assay to expression values obtained by qPCR of the luciferase gene insert (TH-Luc) or the endogenous TH gene (TH11-12). We observed that the luciferase readings of Day 18 plated DA neuron progenitors (following 10-days differentiation) corresponded to the TH expression by qPCR of both TH-Luc and TH11-12, as shown (**Figure 2A-B)**.

**Figure 2.**
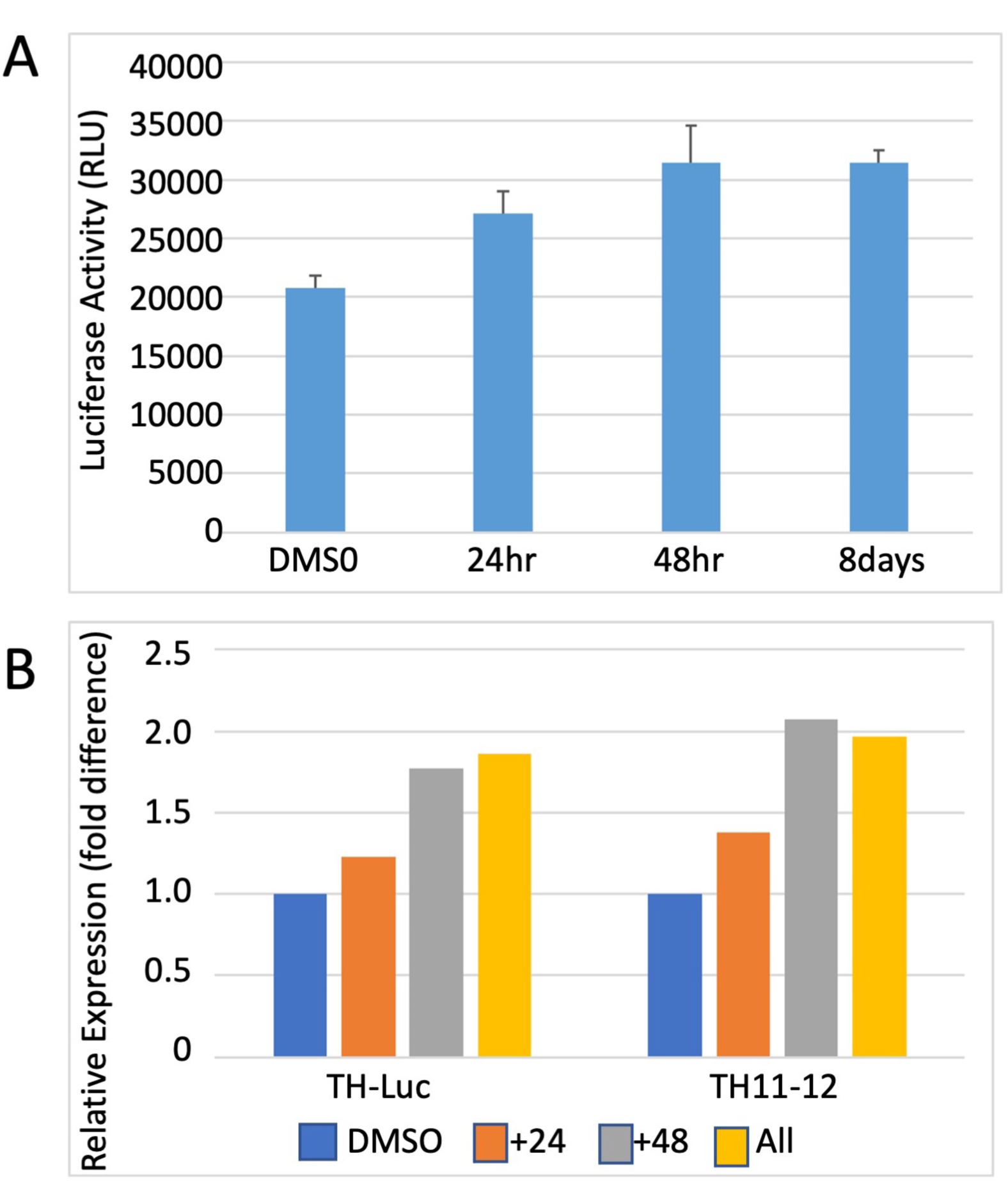
Tyrosine hydroxylase (TH) reporter activity. **(A)** TH expression by luciferase assay. Day 18 DA neuron progenitors were cultured for 10 days with DMSO only (Control) or treated with SGC0946 for the first 24 hr, 48 hr and 8 days. Luciferase readings were done at the end of the 10-day differentiation. **(B)** TH expression by qPCR of the luciferase gene (TH-Luc) or the endogenous TH gene (TH11-12).

Finally, we compared gene expression of a marker of total neurons (TUBB3 (BetaIII Tubulin)) to markers that are reported to be specific to DA neurons (Dopamine Transporter (DAT), GIRK 2 and NURR1). As shown in **Figure 3A**, there is no difference between control and treated total neurons following either single or continuous treatment with SGC0946. In contrast, markers specific to DA neurons (DAT, GIRK2 and to a lesser extent, NURR1) demonstrated an increase in expression with treatment with SGC0946 relative to DMSO Only control as shown in Figure 3B. A longer treatment of SGC0946 increases the expression of DAT, while the expression of GIRK2 and NURR1 can only benefit from a shorter period of SGC0946 treatment during the first 24 or 48 hours of DA neuron differentiation.

**Figure 3.**
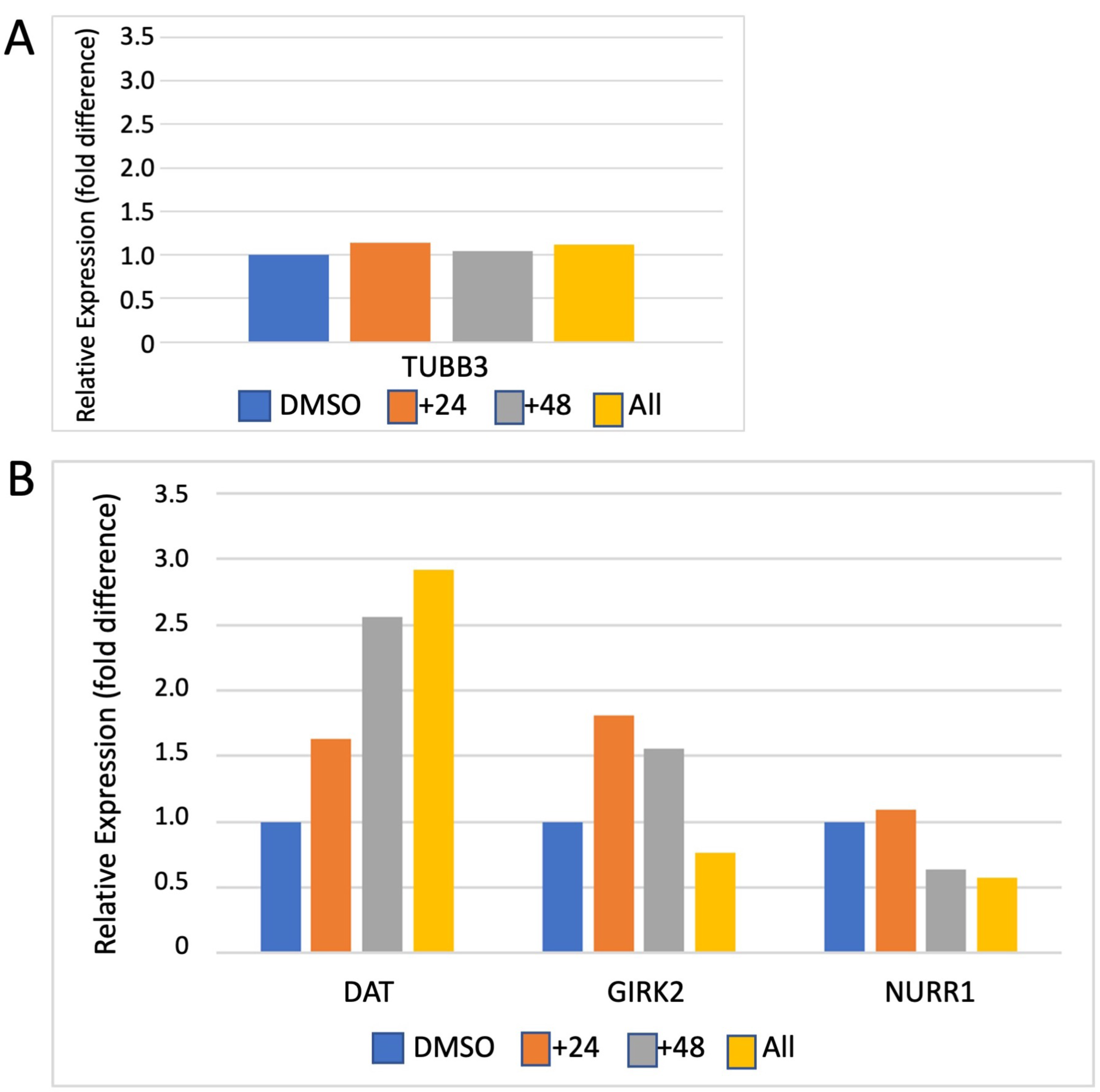
Gene Expression by qPCR. **(A)** SGC0946 treatment does not increase the number of total neurons as measured by expression of TUBB3 (BetaIII Tubulin). **(B)** SGC0946 treatment potentially increases the number of dopaminergic neurons as measured by the expression of TH and Dopamine Transporter (DAT). Data suggests that a short treatment is preferred for the midbrain DA neurons as measured by the midbrain DA neuron marker GIRK 2. NURR1 is a widely expressed marker on neurons, microglia and other cell types.

## Discussion

Over the last decade or more, diverse screening strategies have been proposed for identification of new pharmaceutical candidates to treat PD (Barbuti et al., 2020, Hideshima et al., 2022, Leah et al., 2021, Schikora et al., 2021, Aldewachi et al., 2021, d’Amora and Giordani, 2018, Dawson et al., 2019, Smith et al., 2017, Stewart, 2014, Lavecchia and Giovanni, 2013). Strategies include high-content screening to generate single-cell gene-corrected patient-derived pluripotent stem cell clones useful for further exploration of excess alpha-synuclein with familial PD mutations and identification of perturbed pathways, two-step screening method to identify inhibitors of α-synuclein aggregation, high throughput screens of mitochondrial, neuron or neurite morphology, RPPA (Reverse Phase Protein Arrays) and antibody-based proteomic approaches, and strategies that employ organoids, model organisms and virtual *(in silico)*screening (Barbuti et al., 2020, Hideshima et al., 2022, Leah et al., 2021, Schikora et al., 2021, Aldewachi et al., 2021, d’Amora and Giordani, 2018, Dawson et al., 2019, Smith et al., 2017, Stewart, 2014, Lavecchia and Giovanni, 2013). Given the array of assays that are available for high throughput screening, it is hoped that the use of multiple assays might lead to positive intersections of the most promising treatments for PD, understanding of underlying causes and identification of growth factors and pathways that might be modulated.

Here, we report the use of a human pluripotent stem cell-based system, for discovery of neural toxins and protective factors, in a pilot screen for factors that might enhance differentiation, survival and/or maintenance of DA neurons differentiated *in vitro.* In this pilot screen, we identified SGC0946, an inhibitor of DOT1L (Disruptor Of Telomeric silencing 1-Like) which encodes a widely-conserved histone H3K79 methyltransferase that is able to both activate and repress gene transcription. Our results indicate that SGC0946 increased reporter luciferase activity with a single treatment at 8-hours post-plating being equivalent to continuous treatment. Moreover, data suggested that the total number of neurons differentiated in the assays was comparable from experiment to experiment under different SGC0946 treatments over time. In contrast, data suggested that the survival and/or maintenance of DA neurons might be specifically enhanced by SGC0946 treatment. These results provide evidence in support of other reports that indicate inhibition of DOT1L may play an important role in maintenance and survival of neural progenitor cells (NPCs) and their lineage-specific differentiation (Ferrari et al., 2020, Gray de Cristoforis et al., 2020).

Previously we performed a ChIP-sequencing study and characterized dopamine neuron transcriptional and epigenetic programs including the global binding profiles of H3K27ac, H3K4me1, and 5-hydroxymethylcytosine [5hmC]) at four different stages of development(Xia et al., 2017). We demonstrated the dynamic pattern of epigenetic modification during DA neuron differentiation and maturation. The discovery of the linkage between DOT1L and DA neurons in our compound screen here suggests that H3K79me might also play a key regulatory role during DA development.

As noted in previous primary publications and in a comprehensive review, DOT1-Like (DOT1L) is the sole methyltransferase of histone H3K79 (Wong et al., 2015, Ferrari et al., 2020, Gray de Cristoforis et al., 2020, Wille and Sridharan, 2022). DOT1L can orchestrate mono-di- or tri-methylation that is correlated with actively transcribing genes (mono- and di-methylation) or repression (tri-methylation). Moreover, DOT1L has been recognized as a promising epigenetic target for solid tumors and has been extensively studied in MLL (multi-lineage leukemia) genomic rearrangements and sporadic fusion proteins that may drive errors in the localization of H3K79 methylation and subsequent oncogenesis (Alexandrova et al., 2022). More recently, studies have demonstrated that loss of function of DOT1L is accompanied by perturbations in development of somatic lineages and in reprogramming *in vitro* (Wille and Sridharan, 2022). Thus, the evidence suggests that DOT1L is broadly required for differentiation, that reduced DOT1L activity is concomitant with increased developmental potential and that loss of DOT1L activity results in more upregulated than downregulated genes. DOT1L also participates in various epigenetic networks that are both cell type and developmental stage specific and may play an important role in maintenance and survival of neural progenitor cells (NPCs) and their lineage-specific differentiation (Ferrari et al., 2020, Gray de Cristoforis et al., 2020). Nonetheless, to our knowledge, this is the first association of DOT1L activity and formation, differentiation and/or maintenance of DA neurons. Clearly, further exploration of this association is merited.

## Author Contributions

JC, JoC, and RRP all contributed to the research described in this manuscript: JC, JoC and RRP each contributed to the experimental design, analysis of data and interpretation of the results. JC prepared the manuscript figures; JoC outlined the methods and materials section and RRP prepared the final manuscript for publication.

## Acknowledgments

We thank members of the Reijo Pera laboratory and the McLaughlin Research Institute, past and present, for their helpful comments and review of this work.

## References

Albin, R., Young, A. & Penney, J. 1995. The functional anatomy of disorders of the basal ganglia. Trends Neurosci, 18, 63–4.

Aldewachi, H., Al-Zidan, R., Conner, M. & Salman, M. 2021. High-throughput screening platforms in the discovery of novel drugs for neurodegenerative diseases. Bioengineering (Basel), 8, 30.

Alexandrova, E., Salvati, A., Pecoraro, G., Lamberti, J., Melone, V., Sellitto, A., Rizzo, F., Giurato, G., Tarallo, R., Nassa, G., Weisz, A. 2022. Histone methyltransferase DOT1L as a promising epigenetic target for treatment of solid tumors. Front Genet, 13, 864612.

Bademci, G., Vance, J. & Wang, L. 2012. Tyrosine hydroxylase gene: Another piece of the genetic puzzle of Parkinson’s Disease. CNS Neurol Disord Drug Targets, 11, 469–81.

Barbuti, P., Antony, P., Santos, B., Massart, F., Cruciani, G., Dording, C., Arias, J., Schwamborn, J., Krüger, R. 2020. Using high-content screening to generate single-cell gene-corrected patient-derived ips clones reveals excess alpha-synuclein with familial parkinson’s disease point mutation A30P. Cells, 9, 2065.

Behl, T., Kaur, I., Sehgal, A., Singh, S., Sharma, N., Chigurupati, S., Felemban, S.G., Alsubayiel, A.M., Iqbal, M.S., Bhatia, S., Al-Harrasi, A., Bungau, S., Mostafavi, E. 2022. “Cutting the Mustard” with induced pluripotent stem cells: An overview and applications in healthcare paradigm. Stem Cell Rev Rep, Online ahead of print., doi: 10.1007/s12015-022-10390-4.

Bloem, B., Okun, M. & Klein, C. 2021. Parkinson’s disease. Lancet, 397, 2284–2303.

Byers, B., Cord, B., Nguyen, H., Schüle, B., Fenno, L., Lee, P., Deisseroth, K., Langston, J., Reijo Pera, R.A., Palmer, T. 2011. SNCA triplication Parkinson’s patient’s iPSC-derived DA neurons accumlate α-synuclein and are susceptible to oxidative stress. PLoS One, 6, e26159.

Byers, B., Lee, H., Liu, J., Weitz, A., Lin, P., Zhang, P., Shcheglovitov, A., Dolmetsch, R., Reijo Pera, R.A., Lee, J. 2015. Direct in vivo assessment of human stem cell graft-host neural circuits. Neuroimage 114, 328–37.

Chambers, S., Fasano, C., Papapetrou, E., Tomishima, M., Sadelain, M. & Studer, L. 2009. Highly efficient neural conversion of human ES and iPS cells by dual inhibition of SMAD signaling. Nature Biotechnol, 27, 275–80.

GBD 2016 Parkinson’s Disease Collaborators. 2018. Global, regional, and national burden of Parkinson’s disease, 1990-2016: a systematic analysis for the Global Burden of Disease Study 2016. Lancet Neurol, 17, 939–953.

Cui, J., Rothstein, M., Bennett, T., Zhang, P., Xia, N. & Reijo Pera, R.A. 2016. Quantification of dopaminergic neuron differentiation and neurotoxicity via a genetic reporter. Sci Rep, 6, 25181.

D’amora, M. & Giordani, S. 2018. The utility of zebrafish as a model for screening developmental neurotoxicity. Front Neurosci, 12, 976.

Dawson, J., Warchal, S. & Carragher, N. 2019. Drug screening platforms and RPPA. Adv Exp Med Biol, 1188, 203–226.

Doi, D., Samata, B., Katsukawa, M., Kikuchi, T., Morizane, A., Ono, Y., Sekiguchi, K., Nakagawa, M., Parmar, M., Takahashi, J. 2014. Isolation of human induced pluripotent stem cell-derived dopaminergic progenitors by cell sorting for successful transplantation. Stem Cell Rep, 2, 337–50.

Ferrari, F., Arrigoni, L., Franz, H., Izzo, A., Butenko, L., Trompouki, E., Vogel, T., Manke, T. 2020. DOT1L-mediated murine neuronal differentiation associates with H3K79me2 accumulation and preserves SOX2-enhancer accessibility. Nature Comm, 11, 5200.

Gonzalez, R., Garitaonandia, I., Abramihina, T., Wambua, G., Ostrowska, A., Brock, M., Noskov, A., Boscolo, F.S., Craw, J.S., Laurent, L.C., Snyder, E.Y., Semechkin, R. 2013. Deriving dopaminergic neurons for clinical use. A practical approach. Sci Rep, 3, 1468.

Gray De Cristoforis, A., Ferrari, F., Clotman, F. & Vogel, T. 2020. Differentiation and localization of interneurons in the developing spinal cord depends on DOT1L expression. Mol Brain, 13, 85.

Grealish, S., Diguet, E., Kirkeby, A., Mattsson, B., Heuer, A., Bramoulle, Y., Van Camp, N., Perrier, A.L., Hantraye, P., Björklund, A., Parmar, M. 2014. Human ESC-derived dopamine neurons show similar preclinical efficacy and potency to fetal neurons when grafted in a rat model of Parkinson’s disease. Cell Stem Cell, 15, 653–65.

Hideshima, M., Kimura, Y., Aguirre, C., Kakuda, K., Takeuchi, T., Choong, C.-J., Doi, J., Nabekura, K., Yamaguchi, K., Nakajima, K., Baba, K., Nagano, S., Goto, Y., Goto, Y., Nagai, Y., Mochizuki, H., Ikenaka, K. 2022. Two-step screening method to identify α-synuclein aggregation inhibitors for Parkinson’s disease. Sci Rep, 12, 351.

Hoban, D., Shrigley, S., Mattsson, B., Breger, L., Jarl, U., Cardoso, T., Wahlestedt, J.N., Luk, K.C., Björklund, A., Parmar, M. 2020. Impact of alpha-synuclein pathology on transplanted hESC-derived dopaminergic neurons in a humanized alpha-synuclein rat model of PD. Proc Natl Acad Sci 117, 15209–15220.

Jiang, H., Ren, Y., Yuen, E., Zhong, P., Ghaedi, M., Hu, Z., Azabdaftari, G., Nakaso, K., Yan, Z., Feng, J. 2012. Parkin controls dopamine utilization in human midbrain dopaminergic neurons derived from induced pluripotent stem cells. Nature Comm, 3, 668.

Kedariti, M., Frattini, E., Baden, P., Cogo, S., Civiero, L., Ziviani, E., Zilio, G., Bertoli, F., Aureli, M., Kaganovich, A., Cookson, M.R., Stefanis, L., Surface, M., Deleidi, M., Di Fonzo, A., Alcalay, R.N., Rideout, H., Greggio, E., Plotegher, N. 2022. LRRK2 kinase activity regulates GCase level and enzymatic activity differently depending on cell type in Parkinson’s disease. NPJ Parkinsons Dis., 8, 92.

Kriks, S., Shim, J.-W., Piao, J., Ganat, Y., Wakeman, D., Xie, Z., Carrillo-Reid, L., Auyeung, G., Antonacci, C., Buch, A., Yang, L., Beal, M.F., Surmeier, D.J., Kordower, J.H., Tabar, V., Studer, L. 2011. Dopamine neurons derived from human ES cells efficiently engraft in animal models of Parkinson’s disease. Nature, 480, 547–51.

Lavecchia, A. & Giovanni, C.D. 2013. Virtual screening strategies in drug discovery: a critical review. Curr Med Chem, 20, 2839–60.

Leah, T., Vazquez-Villaseñor, I., Ferraiuolo, L., Wharton, S. & Mortiboys, H. 2021. A Parkinson’s disease-relevant mitochondrial and neuronal morphology high-throughput screening assay in LUHMES cells. Bio Protoc, 11, e3881.

Schikora, J., Kiwatrowski, N., Förster, N., Selbach, L., Ostendorf, F., Pallapies, F., Hasse, B., Metzdorf, J., Gold, R., Mosig, A., Tönges, L. 2021. A propagated skeleton approach to high throughput screening of neurite outgrowth for in vitro Parkinson’s disease modeling. Cells, 10, 931.

Singh, D., Hammachi, F. & Kunath, T. 2017. Modeling Parkinson’s disease with induced pluripotent stem cells harboring α-synuclein mutations. Brain Pathol, 27, 545–551.

Smith, A., Macadangdang, J., Leung, W., Laflamme, M. & Kim, D. 2017. Human iPSC-derived cardiomyocytes and tissue engineering strategies for disease modeling and drug screening. Biotechnol Adv, 35, 77–94.

Stern, S., et al., 2022. Reduced synaptic activity and dysregulated extracellular matrix pathways in midbrain neurons from Parkinson’s disease patients. NPJ Parkinsons Dis, 8, 103.

Stewart, M. 2014. Pluripotency and targeted reprogramming: strategies, disease modeling and drug screening. Curr Drug Deliv, 11, 592–604.

Swistowski, A., Peng, J., Liu, Q., Mali, P., Rao, M., Cheng, L. & Zeng, X. 2010. Efficient generation of functional dopaminergic neurons from human induced pluripotent stem cells under defined conditions. Stem Cells, 28, 1893–1904.

Takahashi, K., Tanabe, K., Ohnuki, M., Narita, M., Ichisaka, T., Tomoda, K. & Yamanaka, S. 2007. Induction of pluripotent stem cells from adult human fibroblasts by defined factors. Cell, 131, 861–7.

Takahashi, K. & Yamanaka, S. 2006. Induction of pluripotent stem cells from mouse embryonic and adult fibroblast cultures by defined factors. Cell, 126, 663–76.

Thomson, J., Itskovitz-Eldor, J., Shapiro, S., Waknitz, M., Swiergiel, J., Marshall, V. & Jones, J. 1998. Embryonic stem cell lines derived from human blastocysts. Science, 282, 1145–7.

Wille, C.K. & Sridharan, R. 2022. Connecting the DOTs on Cell Identity. Front Cell Dev Biol, 10, 906713.

Wong, M., Polly, P. & Liu, T. 2015. The histone methyltransferase DOT1L: regulatory functions and a cancer therapy target. Am J Cancer Res, 5, 2823–2838.

Xia, N., Fang, F., Zhang, P., Cui, J., Tep-Cullison, C., Hamerley, T., Lee, H.J., Palmer, T., Bothner, B., Lee, J.H. Reijo Pera, R.A. 2017. Characterization of dopaminergic neurons derived from a tyrosine hydroxylase reporter stem cell line. Cell Reports, 18, 2533–46.

Yang, D., Zhang, Z., Oldenburg, M., Ayala, M. & Zhang, S.-C. 2008. Human embryonic stem cell-derived dopaminergic neurons reverse functional deficit in parkinsonian rats. Stem Cells 26, 55-63. Stem Cells 26, 55-63., 26, 55–63.

Zhang, P., Xia, N. & Reijo Pera, R.A. 2014. Directed dopaminergic neuron differentiation from human pluripotent stem cells. J Vis Exp, 91, 51737.

